# Do electromagnetic fields from subsea power cables effect elasmobranch behaviour? A risk-based approach for the Dutch Continental Shelf

**DOI:** 10.1101/2023.12.01.569531

**Authors:** Annemiek Hermans, Hendrik V. Winter, Andrew B. Gill, Albertinka J. Murk

## Abstract

Subsea power cables cause electromagnetic fields (EMFs) into the marine environment. Elasmobranchs (rays, skates, sharks) are particularly sensitive to EMFs as they use electromagnetic-receptive sensory systems for orientation, navigation and locating conspecifics or buried prey. Cables may intersect with egg laying sites, foraging habitat and migration routes of elasmobranchs and the effects of encountering EMFs on species of elasmobranchs are largely unknown. Demonstrated behavioural effects are attraction, disturbance and indifference, depending on EMF characteristics, exposed life stage, exposure level and duration. We estimated exposure levels of elasmobranchs to subsea cable EMFs, based on modelled magnetic fields in the Dutch Continental Shelf and compared these to reported elasmobranch sensory sensitivity ranges and experimental effect levels. We conclude that the risk from subsea power cables has a large uncertainty and varies per life stage and species ecology. Based on estimated no-observed effect levels (from 10^-3^ to 10^-1^ µT) we discuss what will probably be the most affected species and life stage for six common benthic elasmobranchs in the Southern North Sea. We identify critical knowledge gaps for reducing the uncertainty in the risk assessments for EMFs effects on elasmobranchs.

## 1. Introduction (1 A4)

The transition from fossil fuel-based energy to renewable energy in the marine environment is considered crucial to meet climate change ambitions and initiatives in many countries [1]. The European Union has set-out targets for an installed capacity of at least 130 GW by 2030 and 1,300 GW by 2050 [2]. The Southern North sea is well suited for the generation of power from offshore wind because it is shallow and average wind speeds are high [3]. Offshore Wind Farms (OWFs) are planned and realised at an increasing rate, covering ever larger marine areas located increasingly further from shore [4]–[6].

The North Sea marine environment is already heavily impacted by human activities, and OWFs represents an additional and large-scale activity that will influence marine life [7]. The effects of scour protection functioning as artificial reefs [8]–[10], exclusion of bottom trawling [11], collision risks for birds [11]–[13] and underwater sound risks for marine mammals [14] are relatively well studied. However few studies exist on the effects of sediment resuspension [15], destratification [15], [16] and marine fauna exposure to EMFs [17]. Different types of subsea power cables (SPC) emit EMFs, including infield cables between turbines, export cables that transmit electricity to shore and interconnector cables that enable power exchange between countries [16], [19]–[22]. SPC produce a locally changed electric and magnetic environment.

Many marine taxa are regarded as magneto-sensitive, whilst some organisms are electrosensitive and a select group, in particular elasmobranchs, are both [18], [19]. Magneto-sensitivity is mainly assumed to be used for long-distance navigation or orientation [20]–[23], which leads to the prediction that the EMFs from SPC export cables that transport electricity to the shore could interfere with migration along the coast. During migration adults will encounter multiple cables of varying EMF strengths which might cause delays as has been shown in migrating eel [24]. Or individuals might show (slight) deviation of course as seen in species of salmonids [25]. A multitude of species respond to magnetic cues and are sensitive to the direction, magnitude or/and inclination of the Earth’s geomagnetic field [26]–[28]. The variations in the geomagnetic field are as low as 0.002-0.005 µT/km between the equator and the poles in relation to a total field strength of between 25 to 70 µT [28], therefore the magneto-sensitivity of species using the geomagnetic field is believed to be in the nano tesla range [25], [26], [29], [32], [28] [36]. Electro-sensitivity is used to find prey [30]–[32], avoid predation [33], [34], and find conspecifics / mates [33]–[35]. Elasmobranchs can detect electric fields as low as 5 (nV/cm) [30].

Benthic elasmobranchs are highly electro- and magneto-sensitive [36] and as a taxonomic group they are already greatly impacted by anthropogenic stressors, such as overfishing, bottom trawling rendering them as bycatch and destroying reef structures needed for reproduction [37], [38]. These fish perform an critical role in the food web linking different trophic levels together and controlling prey populations [39]. As slow growing, often oviparous species with a late sexual maturity, elasmobranch population success is very vulnerable to bottom trawling fisheries and is therefore expected to benefit from the exclusion of these fishing techniques in OWF [37], [40]. Conversely, their benthic ecology places them in direct contact with the extensive (and growing) network of subsea cables emitting EMF. Embryos of oviparous species may be continuously exposed to EMF as export cable routes transverse egg laying sites. The active introduction or recovery of biogenic reefs in OWF might also provide places for elasmobranchs to lay their eggs [40]. Potential effects from EMFs that have been suggested for benthic elasmobranchs, range from [1] disturbance of embryogenic development, [2] behavioural changes such as attraction or avoidance [23] [24], which could lead to disrupted predator-prey relations and/or relations between conspecifics or changes in habitat use and [3] impact on migratory behaviour [42].

Although there is not much known about the effects of EMF on benthic elasmobranchs, their high electro- and magneto-sensitivity and vulnerable life history stages raises concerns, some of which have been expressed by governments [43], NGOs [43] and fisheries organisations [44]. These uncertainties might delay the permitting process [45] or lead to a precautionary approach and the requirement for mitigation measures that are costly and of uncertain added value. A science-based Ecological Risk Assessment (ERA) [46] can support better informed discussions and aid in decision making whether and where mitigation is needed to reduce potential adverse effects [47]. An ERA can also serve to indicate which knowledge gaps should be filled with priority to answer remaining central questions [48]–[50].

This study follows an ERA approach (Figure 1) to assess the risk of EMF emitted by current and planned SPC in the Dutch Continental Shelf in the Southern North Sea for benthic elasmobranchs. The ERA was performed through the following steps:

1. | **Hazard identification:** what is known about the nature of EMFs from SPC, and what indications exist for behavioural responses in benthic elasmobranchs from field observations and experimental studies?
2. | **Exposure quantification:** what EMFs levels have been reported and are to be expected based on modelling from SPC characteristic on the Dutch Continental Shelf?
3. | **Hazard quantification:** what are the (lowest) EMFs exposure levels at which behavioural effects on benthic elasmobranchs occur in exposure studies with different life stages?
4. | **Risk assessment:** comparing highest EMFs exposure levels to lowest effects levels for altered benthic elasmobranch behaviour per life stage in the Dutch North Sea resulting in risk levels.

**Figure 1.**
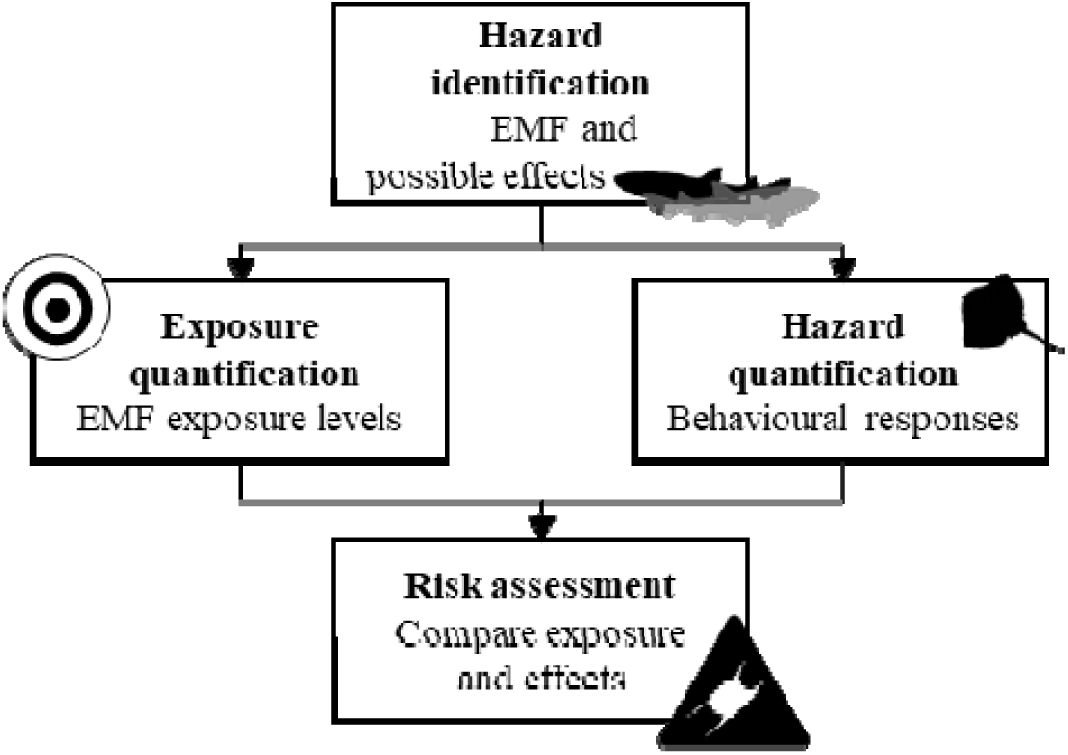
Ecological Risk Assessment approach to determine the risk for behavioral effects of electromagnetic fields (EMFs) resulting from subsea power cables (SPC) on benthic elasmobranchs on the Dutch continental shelf.

The ERA is followed by a discussion in which the results from the risk assessment are related to the different specific traits of the benthic elasmobranchs leading to an estimation of the potentially most affected species/life stage, critical knowledge gaps and the need for appropriate mitigation or retirement of the risk.

## 2. Hazard identification

The hazard identification step evaluated if EMFs may have the potential to cause harm to benthic elasmobranchs. Irrespective of the dose, an overview is provided of the potential risk posed by exposure to EMF and what the characteristics of the EMF are. Below an overview is provided of the nature, sources and potential effects reported by EMFs on benthic elasmobranchs.

### 2.1. EMFs characteristics

The characteristics of the EMFs depend on whether the cable is transporting Alternating Current (AC) or Direct Current (DC). EMFs consists of an electric (E-field), magnetic (B-field) and induced electric field (iE-field). As an industry standard for both AC and DC cables, the direct E-field is contained by cable armouring in a perfectly grounded cable, but the B-field is emitted into the environment and any water movement generates an induced electric or iE field [49]. Furthermore, AC cables also create iE fields owing to the asymmetric cycling of the electrical current in the multiple cable cores. The extent of the magnetic field can vary in relative importance of the directional components (x,y,z plane) [47]. As the magnetic component is a sum of these directional components it therefore does not only have a strength but also a direction [51]. Factors influencing the intensity and direction include amount of power transported, cable design and component geometries, burial depth and cable orientation [49], [52]. EMFs are linearly related to the amount of power transmitted through the SPC which, in the case of OWFs, depends on the available wind. With AC the frequency depends on the local standard, i.e. 50 Hz in most of Eurasia, Africa and Oceania and 60 Hz in North-America. In addition, the form of the cable component rotations (circle or ellipse shaped) can also vary [43]. The vectoral magnetic components interact with other, more variable, natural EMF sources, such as solar radiation (see Nyqvist et al 2020 for an extensive overview of natural EMF sources [28]). iE fields resulting from water movement through a magnetic field, for example tidal currents or water displacement by animal movement. The pattern and magnitude of the resulting iE field depends on the direction and flow of seawater [28], [53] as well as the fish’s size, shape, orientation, and the speed and direction of its movement [28], [54].

### 2.2. Magnetic fields and induced electric fields

The way an organism experiences a specific EMF, and thus the biological relevance, will depend on the species and its life stage [55]. Both the magnetic field and the iE field can serve as potentially important behavioural cues [28], [56]. When an animal is moving through a magnetic field, it simultaneously experiences the magnetic and the induced electric field [28]. Due to this biological coupling these fields cannot be reviewed separately [28], [53]. The iE fields are however difficult to predict and measure as they are heavily dependent on many environmental and biological factors. The exposure quantification applied in this study therefore focused on magnetic fields. As 0.002-0.005µT/km is the latitudinal geomagnetic field (GMF) change for Dutch waters, 0.005 µT was assumed as the minimum level relevant for benthic elasmobranchs to be able to use the GMF for long-distance navigation [28]. Following this assumption, in the risk assessment the lower level of the exposure quantification of 0.005 µT was used.

### 2.3. Potential effects of EMF

Previous studies suggest that EMFs levels that SPC emit are too low to induce acute health effects, such as permanent (neuro)physiological effects in adult individuals [28]. Although the signature of anthropogenic EMFs from a SPC might be different than natural EMFs, Kimber et al 2011 [57] have shown that benthic elasmobranchs cannot discriminate between an artificial and natural EMFs. Indeed, a number of experimental studies address how EMFs induce behavioural responses in elasmobranchs and field observations suggest that elasmobranch interact with SPC [58]. The suggested behavioural effects can be categorized in three types [1] disturbance during embryogenic development, [2] behavioural changes of local-dwelling animals and [3] interaction with migratory behaviour. Potential effects for these categories of behaviour are indicated below and shown in Figure 3.

1. | **Disturbance during embryogenic development** Where SPC come to land they often cross estuaries or other coastal areas potentially well suited as elasmobranch egg laying sites or pupping grounds [59], [60]. Benthic catsharks, such as S*cyliorhinus canicula,* are known to deposit eggs in macroalgae in coastal areas and sessile benthic invertebrates (reefs) on offshore grounds [61]. Benthic skates, such as *Raja clavata,* are known to deposit egg cases in sandy shallow waters of the Southern North Sea [62]. On the Dutch Continental Shelf SPC routes cross the Oosterschelde inlet. If the sessile egg cases are deposited within the EMFs of a SPC, an embryo will be exposed to the varying EMF levels during embryogenesis. Embryos have been shown respond to electric stimuli after 1/3 of their development phase [73] with a freeze response, characterised by temporarily ceasing respiration, suggested to prevent detection by a predator [72]. Constantly responding to changing EMF levels might cost metabolic energy, resulting in reduced growth and development and lead to higher yolk consumption, which may lead to a longer development time and/or smaller hatchlings, as has been seen in lobster larvae [65]. Another impact mechanism could be that continuous exposure to EMFs during embryogenesis could exert epigenetic effects, leading to behavioural alterations in adulthood. Individuals could become less sensitive to EMFs, or the opposite could be true, where individuals could become hypersensitive. These effects might result in reduced foraging success later in life, increased predation risk because they became less sensitive to EMF signatures of predators or a change in habitat use. Also, magneto sensitive species, such as turtles, but also elasmobranchs are believed to use geomagnetic imprinting to maintain fidelity to reproduction sites [66]– [68]. Continuously altering EMFs from SPC during embryogenesis might disrupt the imprinting process needed to find the way back to reproduction sites.
2. | **Behavioural changes during local dwelling** The Southern North Sea provides habitat for approximately 18 elasmobranch species of which 12 have a (partly) benthic ecology [61]. Species abundance and distributions vary but at least some habitats overlap with areas that are suitable for OWF development. Behavioural changes induced by in-field SPC could include altered foraging behaviour and success, disruption in predator-prey and/or conspecifics relationships, or changes in habitat use [48], [69]. For example, anecdotal evidence presented by Barry et al (2008) [70] showed 126 *Raja rhina* aggregating around a not powered SPC compared to no individuals during repeated field survey when the SPC was in use. Hutchison et al reported increased foraging behaviour, such as increased swimming distance and speed, with more sharp turns and changing swimming depth in animals exposed to between 0.3 and 14 µT [36].
3. | **Interaction with migratory behaviour** Benthic elasmobranchs as R*aja clavata, Galeorhinus galeus* and *Mustelus asterias* are known to migrate over long distances up to hundreds km per year [71]–[75]. Migrating elasmobranchs have been shown to orient using the inclination, intensity and angle of the magnetic field [23] Therefore, they might be hindered during migration when crossing a SPC if the characteristics of the geomagnetic field are altered by the emitted EMF. This was observed with eels [24], and with salmonids where individuals were slowed down or showed (slight) deviation of course [25]. Complete blockage of migration is not likely given observed migration patterns from tagged elasmobranchs that must have crossed several existing SPCs [73], [74]. Attraction to the SPC could become point of concern, dependant on the duration of the attraction ranging from short (seconds) as shown by Orr (2016) [76] to longer attraction as shown by Barry *et al* 2008 [70]. Cumulative minor migration alterations and delays might have energetic consequences resulting in reduced migration success, missing important ecological timing, reduce overall fitness or increase predation risk when crossing SPC higher in the water column.

**Figure 3.**
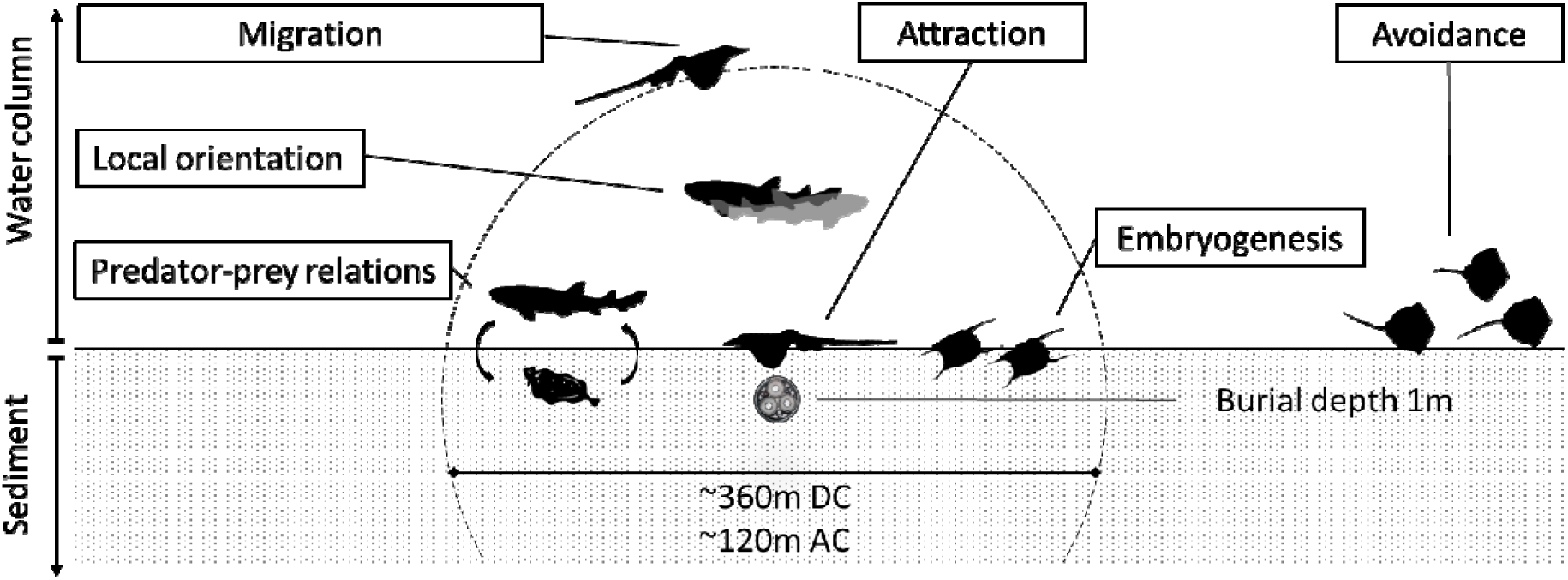
Schematic overview of possible elasmobranch responses exposed to modelled magnetic fields (Figure inspired by Albert et al. 2020). Potential impact range based on a perception level of 0.005 µT, modelled levels for the OWF export cable IJmuidenVer (2GW direct current subsea power cable) and Borssele (700 MW alternating current subsea power cable) transporting maximum amount of power is indicated by dotted line. Note: animals are not to scale.

## 3. Exposure quantification

### 3.1. Measured magnetic field levels on the Dutch Continental Shelf

In order to assess the risk of any stressor, it is important to quantify the exposure that animals may encounter, estimating frequency of occurrence, variability in EMFs and the range of field strength. To be able to estimate the current and for the nearby future predicted EMFs levels as potential exposure of benthic elasmobranchs in the Southern North Sea, a literature review was performed on the measured magnetic field levels over three phase AC and bundled bipolar DC SPC (Table 1). The majority of the measurements were performed at the seabed, where the distance to the SPC depends on the burial depth, generally 1.0-2.0 m deep. The reported magnetic field levels above cables transporting AC or DC range from 0.004 µT to 6.540 µT for AC cables and 0.46 µT to 20.7 µT for DC cables. The magnetic field levels depend on distances to the SPC, environmental conditions (flow, temperature etc.) and power transported through the SPC at the time of measurement, and could not be measured during high wind conditions for safety reasons. Therefore, the scenario for high intensity EMFs can only be modelled. For assessment of the spatial range in exposures benthic elasmobranchs might encounter, the spatial and temporal characteristics of EMF should be taken into consideration and can be approximated with modelling [47].

**Table 1.** Magnetic field levels measured above alternating current (AC) 3-phase and bundled bipolar direct current (DC) export and interconnector cables. The values noted in the table are the maximum levels recorded during a measurement. The power transported through the cable at the moment of measurement was not provided in the listed studies.

| Alternating Current (AC) Cables - 3-phase |  |  |  |  |
| --- | --- | --- | --- | --- |
| Cable specifications | Maximum levels (μT) | Distance from the cable (m) | Reference |  |
| 34 kV; 108 MW | 0.008 - 0.020 | 1.5 - 2.0 | Snoek et al., 2020 |  |
| 150 kV; 120MW | 0.015 - 0.039 | 1.5 - 2.0 | Snoek et al., 2020 |  |
| 150 kV; 129 MW | 0.004 | 1.5 - 2 | Snoek et al., 2020 |  |
| 50 Hz | 0.004 | 1.0 - 1.5 | Thomsen et al., 2015 |  |
| 50 Hz; 70 A | 0.017 | 15 | Thomsen et al., 2015 |  |
| 36 kV; 265 A | 0.600 | 1.5 | Gill et al., 2009 |  |
| 33 kV; 50 A | 0.050 | 0 | CMACS 2003 |  |
| 11 kV; 60 A | 0.056 | 0 | CMACS 2003 |  |
| 150 kV; 120MW; 436 A | 6.540 | 0.5 | DNV-GL, 2015 cited in Snoek et al., 2020 |  |
| Direct Current (DC) Cables - bipolar |  |  |  |  |
| Cable specifications | Magnetic field levels (μT) | Maximum levels above GMF (μT) | Distance from the cable (m) | Reference |
| 300kV; 330 MW; 0 A | 51.7 | 0.46 | 1.3 | Hutchison et al., 2020 |
| 300kV; 330 MW; 16 A | 51.9 | 0.64 | 1.3 | Hutchison et al., 2020 |
| 300kV; 330 MW; 345 A | 65.6 | 14.3 | 1.3 | Hutchison et al., 2020 |
| 660 MW; 500 kV; 1320 A | 72.0 | 20.7 | 1.3 | Hutchison et al., 2020 |
| 660 MW; 500 kV; 660 A | 56.3 | 4.7 | 1.3 | Hutchison et al., 2020 |

### 3.2. Modelled magnetic field levels for the Dutch Continental Shelf

As a case study we modelled the local magnetic field levels (in µT) of SPC at the seabed of the Dutch Continental Shelf. The model comprises a 24 year period, from the first cable installation in 2006 up to the planned SPC network to connect the allocated OWF by 2030. Telecommunication cables were not included as their EMF levels are believed to be negligible (<0.005 uT) because of low voltage (10 kV) [48], [77], [78]. The magnetic fields of 3-phase (commonly used in AC SPC) and bipole (for DC SPC) cables were calculated using the Biot-Savart law (for a detailed description of the calculations refer to the supplementary materials S1). Cable characteristics were obtained from publicly available documentation and/or provided by the cable owners (for details see supplementary materials Table 1). The burial depth was assumed to be the legally required minimum of -1m in Dutch waters.

As the capacity of the AC SPC cables has increased in the last decades, the modelled magnetic field increased from a maximum of 1.8 µT in the first Dutch OWF *Windpark Egmond aan Zee* (OWEZ) which has been operational since 2007 to a maximum of 13.9 µT for the recently build OWF Borssele (Figure 2a). This increase in magnetic field strength also can be observed for DC cables where the capacity and thus output has quadrupled in the same time span from 32.7 µT in NorNed (built in 2008) to 122.7 µT for IJmuiden Ver planned for 2028 (Figure 2b).

**Figure 2.**
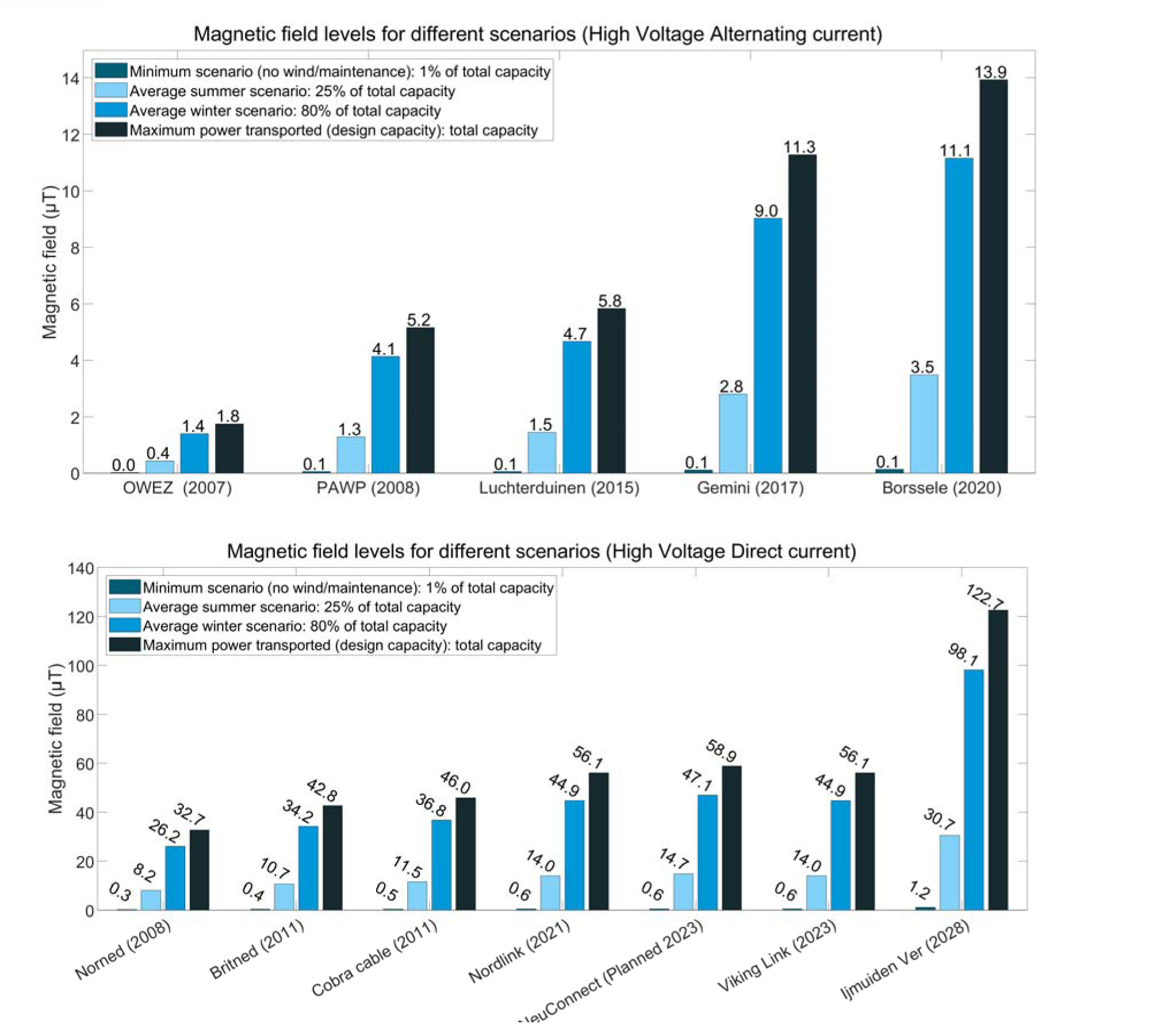
Modelling results of produced magnetic field levels from existing and planned SPCs up to 2030 of (a) from Offshore Wind Farm export and (b) interconnector cables on the Dutch Continental Shelf. Note that the Borssele cable also represents the Hollandse Kust Zuid, Noord and West, and Ten Noorden van de Wadden cables and the Ijmuiden Ver cable also represents the Nederwiek cables respectively which are planned but not shown as the cable specification and thus the modelled values are the same. The scenarios are defined as a percentage of the design power (DP) minimum (1% of DP, only maintenance power levels), summer (25% of DP), winter (80% of DP) and maximum (100% of DP).

As the amount of transported energy and therefore the magnetic field levels above SPC depends on the wind conditions, we modelled seasonal scenarios to provide a realistic exposure range. We assumed the production of an average of 25% of the maximum (designed) power in summer and 80% of the designed power in winter (Figure 2). The percentages used for the scenarios are based on information supplied by the grid operator. For example, a commonly used 700 MW 3-phase AC (Borssele) summer scenario resulted in an average magnetic field of 3.5 µT and the winter scenario 11.1 µT. Without wind, the maintenance power level including the reactive power results was modelled at 0.1 µT and the maximum EMF that the cable can produce is 13.9 µT. The modelled magnetic field levels for planned DC cables are almost an order of magnitude higher than for AC cables (Figure 2). Unless OWF generated wind will be converted into e.g. hydrogen at sea, future scenarios (2030-2050) will include more cables with higher capacities transporting electricity over longer distances [4] which will further increase EMFs exposure levels [47].

### 3.3. 6% of Dutch Continental Shelf covered with EMF detectable by elasmobranchs by 2030

The magnetic field levels exponentially decrease with distance from the SPC. Based on the lowest perception level of 0.005 µT the affected area of SPC from export and intercontinental cables considered in this study totals over 850 km^2^. This is approximately 1.5 % of the Dutch Continental Shelf (∼57,800 m^2^). The potential impact range of 0.005 µT contour differs from roughly 40 metres in diameter for the OWEZ cable to 360 metres for the IJmuiden Ver SPC (supplementary materials Table 2). It is important to consider that as the EMFs extend in all directions, and the extent can differ horizontally and vertically, and B and iE fields are perpendicular to each other. For this risk assessment it is assumed that this extension is the same in all directions. This means that not only the seabed but also (most of) the water column is influenced by EMF as the water depth is a maximum 50 meter in the DCS. Besides the export cables, the inter-array cables within the OWF also create an EMF. OWF are planned to occupy 2,600 km^2^ in 2030 which is circa 4,5% of the Dutch Continental Shelf. Assuming the EMF from the inter array cables covers the entire OWF, circa 6% of the Dutch Continental Shelf will be under the influence of SPC generated EMF detectable by elasmobranchs by 2030. And, of course, this exposure will not be limited to the Dutch Continental Shelf as in the surrounding North Sea countries comparable OWF developments are taking place, aiming for a total energy production 130 GW by 2030 in the entire North Sea [2].

### 3.4. Rate of encounter

The elasmobranchs habitats will not fully overlap with this 6% of the Dutch Continental Shelf under the influence of EMFs. The species-specific movement ecology and habitat uses will determine encounter rates. For most benthic elasmobranch species these life history characteristics are not yet known. As a first proxy for encounter rate, overlap in spatial occurrence of benthic elasmobranchs and areas under the influence SPC induced EMFs is examined. For this, the number of species caught in the Frisbe and DATRAS fisheries surveys of ICES at a certain location in a period of ∼40 years (data collections methods described by Batsleer et al (2020) [38]) is overlayed by the installed and planned SPC until 2030 (Figure 5). This shows that OWFs overlap habitat of up to four of the studied benthic elasmobranch species. Some export cables (IJmuidenVer and Borssele) transvers areas are used by up to 5 different species for either migration of local-dwelling.

**Figure 5.**
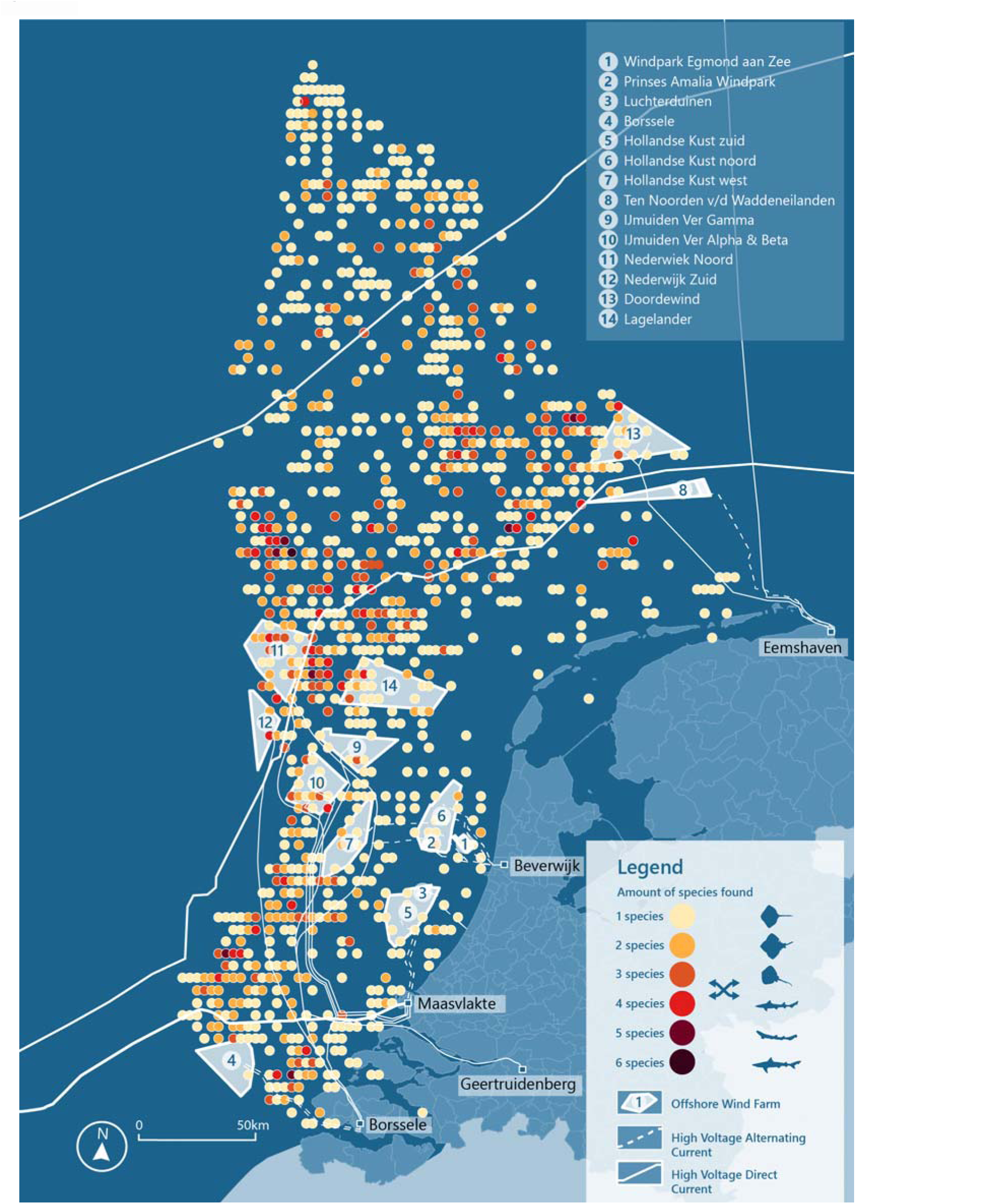
Overview of offshore wind farm export cables and interconnect cables build and planned until 2030 on the Dutch Continental Shelf in relation to the catch data of elasmobranch species. Solid lines indicate direct current cables, dashed lines indicate alternating current cables, data obtained from Informatiehuis Marien Open data viewer. Species richness indicates number of elasmobranch species caught based on Frisbe (1980 – 2019) and DATRAS (ICES) fisheries database. Figure shows overlap between cable routes and habitat use of Elasmobranchs.

### 3.5. Comparing measured and modelled EMF levels

It is only possible to validate the modelled magnetic field levels with measured EMFs levels if key information that influences field strength is available, such as power transported through the cable at the time of measurement, burial depth and cable characteristics [51]. This is mostly lacking, resulting in apparent discrepancies between measured and modelled magnetic field levels [79], [80]. Hutchison et al (2020) reported AC levels of >0.005 µT tens to a hundred of metres from the cable recorded at DC cables [51], which was unexpected. The most likely explanation was interference from the transformation stations. Other measurements report AC magnetic field background values of ± 0.0325 µT, which were higher than the EMF in references areas without SPC [81]. Thomsen et al (2015) described that magnetic field levels measured at Belgium OWF export cables were higher than calculated [79]. On the other hand, Snoek et al. (2020) measured magnetic field levels up to 0.039 µT at the AC ‘PAWP’ SPC located in the Dutch North Sea, transporting power generated by wind levels at 3-4 Bft [81]. Our modelled levels for those conditions, assuming standard burial depth, were >30 times higher (1.3 µT in the comparable wind scenario, summer scenario, 25% of total capacity) (Table 1 and Figure 2). The theoretical transported power does not always reflect the actual transported power and there is an imbalance between the three phases in AC cables that can explain this difference. In addition, the 3-phase cables are helically twisted which reduces the magnetic field level as the fields of the three cores party cancel each other out. This cancellation could not be taken into account in the model, as it would require information on the twist length periodicity (the cores have made a 360° twist around each other), which is currently not provided in cable specifications [82]. For bundled DC cables the orientation in which the cables are placed on/in the seabed (vertical, horizonal or diagonal) influences the interaction with the geomagnetic field. The placement orientation is generally not known and also not included in the existing models. Therefore, measurements are still needed to account for these uncertainties, and modelled values to be able to include EMFs generated at the highest wind levels. The combination of both methods can provide understanding in the critical knowledge gaps needed to be able to quantify EMF exposure levels for benthic elasmobranchs.

## 4. Hazard quantification

To determine the risk of behavioural effects the No-Observed and/or Lowest Observed Effect Levels (N/LOEL) need to be established to be able to compare these to the exposure levels for the risk assessment. The (No-) effect levels will depend on the species/life stage and exposure type/duration and studied endpoint. In this hazard quantification the N/LOEL was established based on the available literature. EMFs can induce behavioural changes in benthic elasmobranchs and based on a literature review of laboratory and mesocosm studies on behavioural responses to EMF exposure, the No- or lowest effect levels were assessed for DC as well as AC induced EMFs (Table 2). In total 9 studies were found, which is very limited for a good hazard quantification. Seven studies were laboratory based and two mesocosm based. When determining the LOEL it is important to consider that dose-response curve is potentially not S-shaped, but maybe show a downward trend or (inverted) U-shape [79]. Lower levels of EMFs might have different effects as they could evoke attraction of a longer duration with less individual variation, which for example may mimic bioelectric fields of prey [57]. This might explain why some studies that include high EMF levels [19], [83] show no effects, whist others with lower levels that are comparable to the levels expected at on the Dutch Continental Shelf, reveal behavioural changes [36], [41].

**Table 2.**
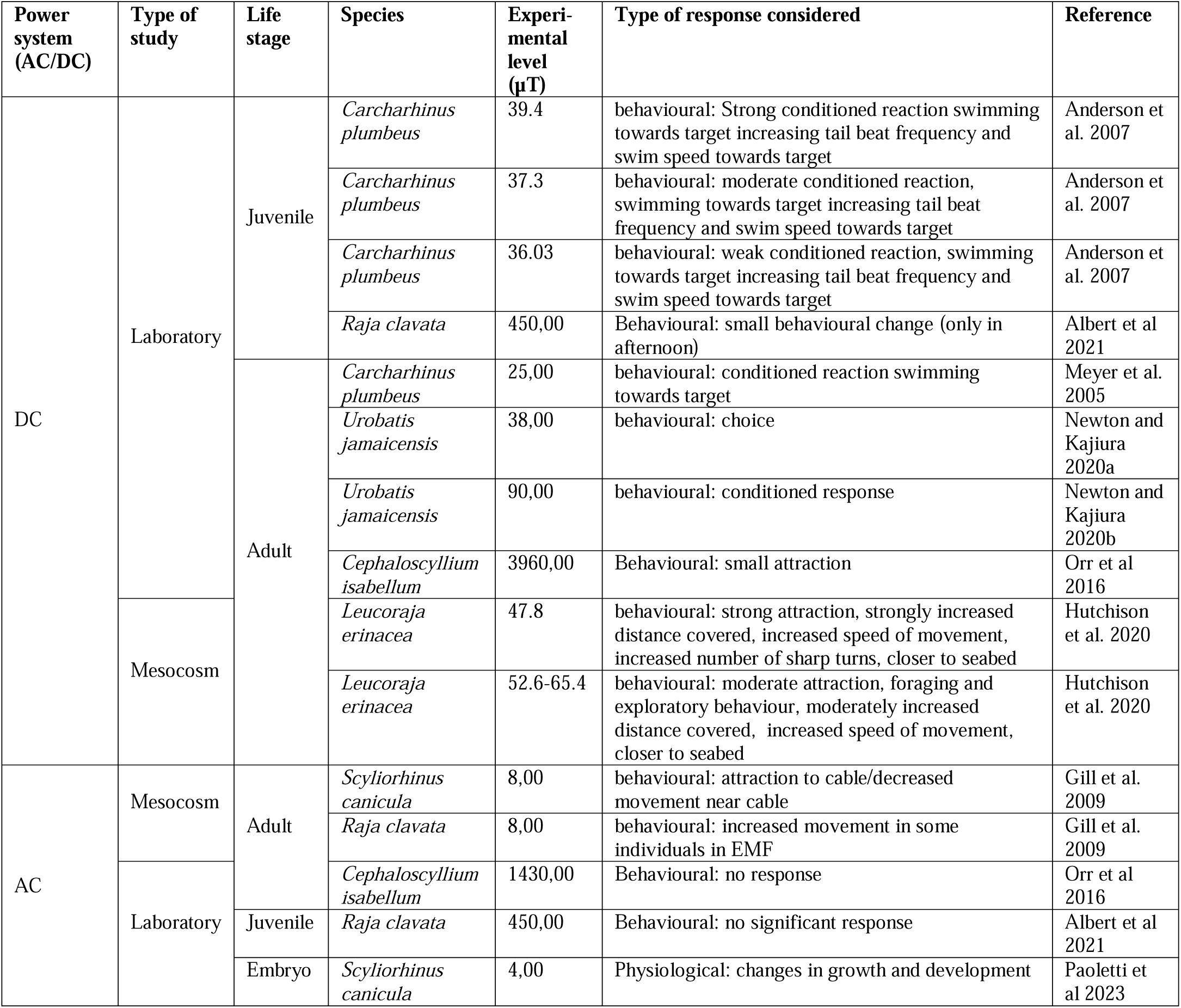
Overview of reported benthic elasmobranch behavioral responses to anthropogenic magnetic field exposure of alternating (AC) and direct current (DC) power transmission systems or changes in the geomagnetic field (GMF)

### 4.1. Reported effects of EMF

The behavioural responses to EMFs exposure that are described literature are summarised in Table 2. The reported effects are clustered for the three behavioural categories distinguished in the hazard quantification: embryonic, local-dwelling and migratory behaviour.

1. | **Disturbance during embryogenic development** There is only one study that looked into the impact of EMFs during the embryonic phase [84]. Paoletti et al found no changes in freezing response, nor ventilation frequency, in *Scyliorhinus canicula* eggs exposed to a constant EMFs levels of 4-7 µT 50 Hz AC during the entire embryogenesis. They also found no changes in stress hormones (1α-hydroxycorticosterone) levels. The authors did find a subtle change in growth rate in the second half of development. The exposed individuals were more advanced in development (smaller yolk sack, larger body size) than the control group at 18 weeks. In the study of Paoletti *et al* the embryos were not followed until hatching so an influence on behaviour after hatching was not determined. The effect on development period, size at hatching and possible developmental alterations could be stronger after longer development. The only available LOEL for AC causing a subtle effect after exposure during 27 weeks is 4 µT. To extrapolate to NOEL, taking into account the relatively short exposure and observation period we apply an extrapolation factor of 10 as is common practice in ecotoxicology, resulting in a LOEL-AC of 0.4 µT. No studies are reported to estimate a N/LOEL for DC exposure. However, it is hypothesized that the effects of DC may be more pronounced compared to 50 Hz AC (AC). This is because DC is considered to be more perceptible and similar to the bioelectric field emitted by a potential predator which causes embryonic stress [85]. We therefore adopt a preliminary NOEL of 0.4 µT for both AC and DC as a rough estimation.
2. | **Behavioural changes during local-dwelling** In a study by Hutchison et al 2020 *Leucoraja erinacea* was exposed in a field-based mesocosm study to a SPC with levels from 0.03 µT DC above the GMF and showed increased swimming distance and speed, more sharp turns and swimming closer to the seabed [36]. In the same study similar but stronger responses were found at higher levels up to 14 µT DC above GMF. These findings are supported by Anderson et al. 2007 who showed a conditioned response to a magnetic field target at the same minimum value of 0.03 µT DC above GMF. in juvenile *Carcharhinus plumbeus*. The response increased with magnetic field strength and stronger responses were noted at 5.3 µT DC above GMF [18]. The LOEL was established at 0.03 µT DC.

The LOEL value at which a behavioural response was recorded for AC was 8µT in another mesocosm experiment with adult *Raja clavata* and S*cyliorhinus canicula* [41]. Here, for *Raja clavata* some increase in movement was seen and for *Scyliorhinus canicula* attraction to the SPC combined with decreased movement near the SPC was observed. At higher levels some attraction of *Cephaloscyllium isabellum* to DC was recorded at 3960µT for DC but not for AC levels at 1430 µT [76]. Albert et al 2021 also worked with high exposure levels (450µT DC) of juvenile *Raja clavata* and found a small increase in movement [83], which was dependent on the time of day. The same experiment with AC did not show any changes in behaviour of juvenile *Raja clavata*. None of the reviewed studies found indications for avoidance behaviour. Avoidance is only known for shark deterrent devices in which extremely high magnetic field levels are applied (i.e. 250 000 µT) [86]–[88]. The lowest LOEL for DC is 8 µT. There are indications of a non-S shaped dose-response curve, suggesting lower EMF levels might result in more effect then higher levels as bioelectrical signals of fish (mates, partners and predators) are much lower [85]. Extrapolation from the LOEL to a NOEL for AC is therefore set at an order of magnitude lower, at 0.8 µT AC.

1. | **Interaction with migratory behaviour** No studies were performed to investigate migratory responses by (benthic) elasmobranchs to EMF from SPC. Based on field observations of delayed migration in eel and salmon [24], [25] and based on the reasoning that the benthic elasmobranchs can navigate differences in the geomagnetic field as low as 0.002 - 0.005 µT we assumed that the NOEL for confusion or diversion caused by migrating over SPC are expected to be around 0.005 µT. It is anticipated that disruptive effects may occur when the levels are increased by a factor of 10, placing the NOEL at 0.05 µT. As the geomagnetic field (DC) is used for migration 50 Hz AC levels are not believed to impact migration and therefore no effect thus no risk for AC is assumed for this endpoint.

## 5. Risk characterization and Assessment

When overlaying the exposure levels presented in literature (Table 2) with the modelled EMFs levels from the exposure quantification (Figure 2) an overlap of the sensory range of benthic elasmobranchs can be observed (Figure 4). This means there is reason to continue with the ERA where the exposure levels are compared with the NOELs to determine whether the risks occur, to be determined per life stage category. In case there is reason for concern, the spatial encounter risk is determined for the different life stage categories and related to the ecological traits of elasmobranch species.

**Figure 4.**
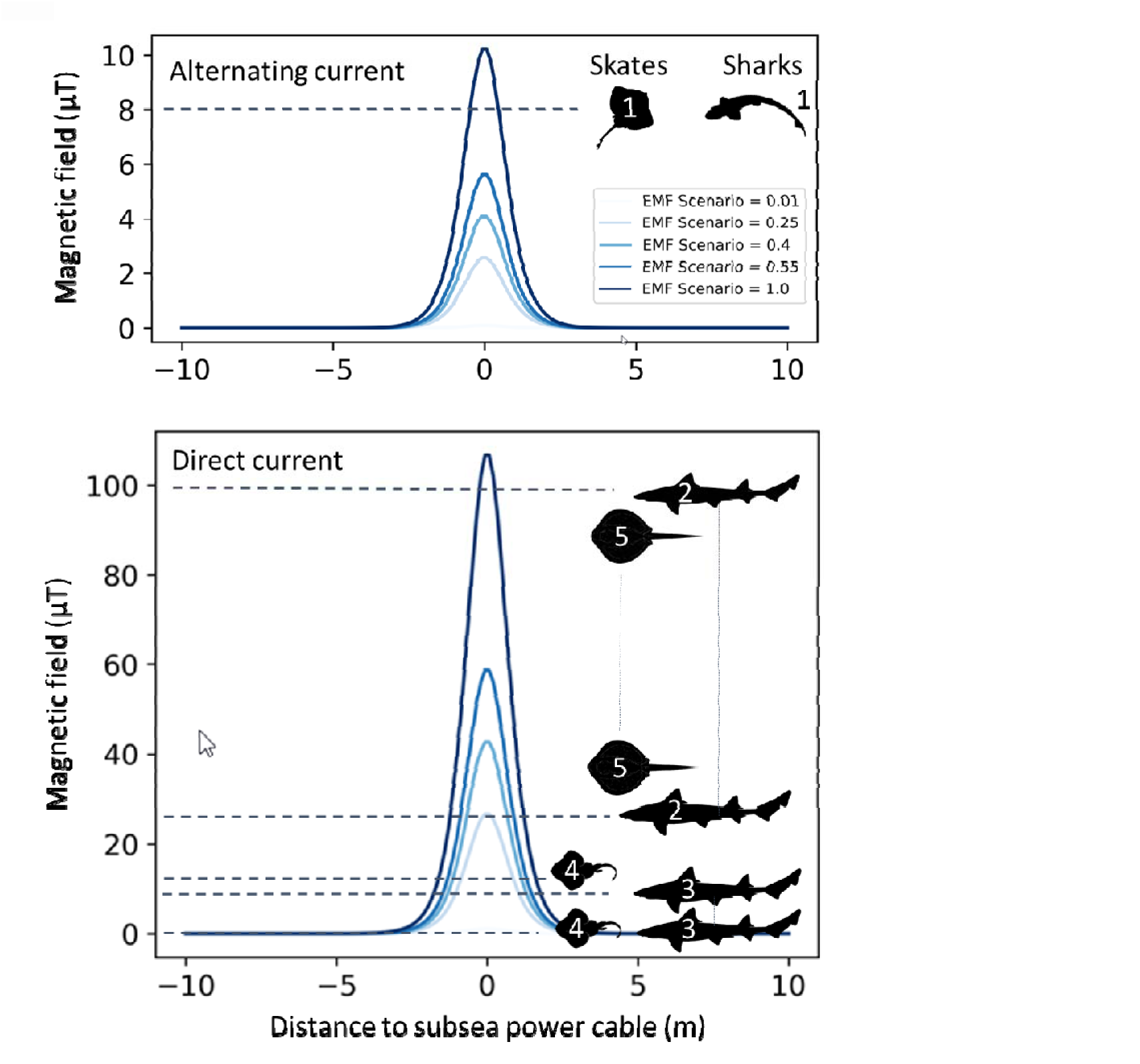
Schematic overview showing overlap of [1] modelled magnetic field levels of alternating current (top) and direct current (bottom) subsea power cables and [2] elasmobranch responses to magnetic stimuli based on mesocosm and laboratory studies (dotted lines). The number in the depicted animals correspond with the studies: [1] Gill et al. 2009: 8 μT, [2] Meyer et al. 2005: 25 to 100 µT, [3] Anderson et al. 2007: 0.03 to 8.00 µT, [4] Hutchison et al 2020: 0.3, 4.0 and 14 µT [5] Newton and Kajiura 2020 38, 45, 90 µT. Note that lower magnetic field levels (nT) are not visible as the y-axis is not logarithmic, but can extent up to 120 (AC) and 360 (DC) meter from the subsea power cable, depending on cable characteristic and power levels transported.

As the maximum magnetic fields cannot be measured, only modelled exposure levels, we used these as input for the risk assessment: max 122.7 µT DC (projected for IJmuiden Ver, 2028 onwards) and 13.9 µT for AC (current for the OWF Borssele), respectively.

The hazard quantification provided an NOEL of 0.4 µT for embryonic development and 0.4 µT and 0.8 µT for DC and AC, respectively, for behavioural changes during local dwelling. For interaction with migratory behaviour an DC NOEL of 0.005 µT for confusion/diversion and 0.05 µT for disruptive effects is adopted.

### 5.1. Risks assessed per effect

1. | **Disturbance during embryogenic development** The maximum EMFs levels elasmobranch embryos are potentially continuously exposed to were 13.9 µT for AC cables and 122.7 µT for DC cables. The exposure levels are highly variable due to the changes in power transported and are likely to vary hourly (small changes) and daily (larger changes). The NOEL for impact on embryonic development is 0.4 µT for both AC and DC. The likelihood of encounter was judged as likely due to the overlap of cable routes and areas that are suitable as egg laying sites which will mean continuous exposure of egg cases when laid within the vicinity of SPCs. The NOEL is two and three orders of magnitude lower than AC and DC exposure levels respectively, which is reason for concern. While experiments are done with constant EMF levels, an extra risk lies in the constant changing EMF level as this may reduce the potential for habituation. As embryo’s especially react to changing EMF levels, constant adjustment might require extra energy which might disturb growth and development.
2. | **Behavioural changes during local-dwelling** Both infield within the OWF and export cables, transporting the power from the transformation station to shore, can be encountered during local-dwelling by benthic elasmobranchs and might cause behavioural effects. However, the chances of encountering an SPC is largest within OWFs as the cable density is higher. In addition, the infield cables form a network like structure of cable strings connect the turbines to the transformation station. As the EMF levels of infield cables are lower than of export cables this does mean there is likely less effect. Specifically with the possibly non-S-shaped dose-response relationship, exposure to lower EMF levels might result in more confusion or disturbance then higher levels. Infield cables are not modelled as part of this ERA but are comparable to the modelled levels for the OWEZ export cable. However, the OWEZ cable has comparable characterises to the majority of the infield cables and was used as a proxy for EMF levels. The maximum magnetic field level based on modelling was 1.8 µT. The NOEL for behavioural changes resulting in (different types of) local behaviour changes was set at 0.4 µT and 0.8 µT for DC and AC, respectively. The likelihood of encounter was judged as possible because circa 6% of the Dutch continental shelf is expected to be under influence of EMF in 2030. Most OWF and SPC routes overlap with habitat of up to 6 species of benthic elasmobranchs (Figure 5). The NOEL was an order of magnitude lower than the exposure levels indicating risk of behavioural changes within OWFs. The main risk lies in long term attraction which could result in wasted energy expenditure or possibly changes in foraging and habitat use of 6% of the Dutch continental shelf.
3. | **Interaction with migratory behaviour** The maximum exposure level for migrating individuals was 122.7 µT for DC cables; specifically DC SPC interact with the geomagnetic field. The NOEL for potential confusion to migratory behaviour was set at 0.005 µT and for disruptive effects at 0.05 µT, while the possibility of exposure was classified as high as SPC routes are perpendicular to the coast (Figure 5) suggesting multiple encounters when migrating over larger distances. An animal migrating from the North to the South along the Dutch coast might encounter up to 31 cables in 2030 (Table S1). When it is traveling longer distances in the Southern North Sea or adjacent seas it might encounter even more cables in Germany, the United Kingdom, Belgium and France. The NOEL for confusion was five orders of magnitude lower, and for disruption it was four times lower than exposure, suggesting a considerable risk and reason for concern for interaction of EMF with migratory behaviour.

## 6. Discussion

In this ERA exposure to EMF from SPC resulted in risks for three different categories of behaviour, [1] disturbance during embryogenesis, [2] behavioural changes during local-dwelling and [3] interaction with migratory behaviour. During embryonic development and migratory behaviour the risks were most pronounced. The number of available studies to base the ERA on is limited, and all those studies were based on a small sample size and/or report large individual variation [23], [24], [62], [63] resulting in a low level of confidence. It is strongly recommended that the N/LOELs reported in this study should be refined. The NOEL for confusion during local dwelling was based on a hypothetically lower sensory range of (benthic) elasmobranchs used for navigation [28]. This level (0.005 µT) might be an under estimation of the lower sensory threshold for navigation, which may mean the actual total spatial impact zone is larger.

### 6.1. Species traits dependent risk estimation and priority for further research

In order to provide an impression of the risks from EMF exposure of benthic elasmobranchs with different life cycle traits, to facilitate determining research priorities, a risk matrix was created to categorise risks ranging from no to high concern. This matrix was based on the probability of the species and life cycle being exposed to EMFs from SPC (y-axis) combined with the severity of the impact (x-axis) [89], [90] for the six most common benthic elasmobranchs on the Dutch Continental Shelf. The resulting risk qualification depends on the biology and life histories of different species, e.g. rays, sharks or skates, being resident or migratory, and oviparous or ovoviviparous/viviparous species (Figure 6).

1. | **Disturbance during embryogenic development** Based on the effects of EMFs on exposed embryos or eggs in other species the severity of impact could range from low (negligible impact), medium (impact on yolk consumption and decreased size) to high (deformities and reduced swimming behaviour). For oviparous species that lay their eggs on sandy substrate such as *Raja clavata* it is known that nursery areas are located in shallow sheltered areas, which are also often used for the landing of SPCs. Species like *Amblyraja radiata* are believed to lay eggs throughout the North Sea, including deeper waters [61]. This wide distribution of eggs indicates a likelihood of overlap between SPC and egg cases. The probability of encounter can range from low to high depending on the actual overlap of cable routes and egg laying sites. *Scyliorhinus canicula* is an example of an oviparous species that uses reef like structures and benthic invertebrates to attach egg cases [91]. SPC on the Dutch Continental shelf are laid in soft sediment areas. Therefore, any exposure might be less likely but the rocks placed on cables crossings, or scour protection of turbines offer potential habitat heterogeneity which might make suitable egg deposition sites. The probability of encounter is therefore judged as low to high. The highest risk can be identified when egg laying sites and SPC routes directly overlap. This would result in continuously exposed egg cases with a potential high severity e.g. physical changes, such as deformities or changed EMF perception. Research should focus on the effects of EMFs during the complete embryogenesis of oviparous elasmobranchs. Of the two power systems used in OWF, DC systems will have a greater spatial extension owing to higher intensities. In addition, DC interacts with the GMF, whereas AC is not expected to and therefore DC research should therefore be prioritized. Conversely, AC could lead to different effects especially in relation to the varying levels (mimicking low wind and high wind days) and potential effects should therefore also be researched. Controlled laboratory experiments would be very suitable to study the effect of EMFs on embryonic development in egg cases. Following a laboratory approach, it is important to choose a model species of which egg cases can be obtained in sufficient numbers, are not classified as endangered and can be reared successfully in captivity as for example *Raja clavata* and *Scyliorhinus canicula*.
2. | **Behavioural changes during local-dwelling.** The local-dwelling phase is not, at present, a priority risk requiring further research. None of the 6 species considered has severity of consequences deemed to be high (Figure 6). Except for common stingray and tope shark, however, it cannot be excluded that EMF exposure could be high. The unknowns in species ecology include the spatial extent of foraging behaviours and local habitat preference which all affect the probability of encountering SPCs. For species that forage closer to the seabed such as *Mustelus asterias* the chances of disruption in predator prey relations are more likely than for species that (also) feed pelagically as *Galeorhinus galeus* [61]. The impact on foraging might be indirectly increased further for species that feed on prey that might also be influenced by EMF as crustaceans [92] as is the case for *Mustelus asterias* but less so for the piscivore *Galeorhinus galeus.* The location (depth and distance to shore) of the OWF also influences the encounter rate as the habitat suitability will differ between OWFs (Figure 5). For example *Raja clavata* is more frequently caught offshore and *Amblyraja radiata* has a more northly orientated distribution range [61]. The effects described in literature include impact on foraging behaviour and short-term attraction, which are reasons for concern but the consequence of severity is considered low to medium. Although there are no studies that suggests avoidance, there is only anecdotal evidence of benthic elasmobranch presence in OWFs and no information to which extent habitats within OWF are used. The main risk lies in OWFs becoming unsuitable habitat for foraging or mating due to the vast network of infield cables. Research could focus on habitat use within OWFs using for example eDNA methods [93]. By establishing behaviourally focused dose-response relationships using varying levels of AC comparable to the EMF levels generated by in-field SPC, valuable insights can be gained regarding the potential disruptions in conspecifics or predator-prey relationships. It is anticipated that these relationships will be most affected, as AC fields are similar to the natural fields produced by fish [30], [94]. Research on behavioural changes in OWFs should focus on the species which are expected to have a long(er) resident time within these parks. Studies of this nature can be done in a laboratory setting or in the field with Baited Remote Underwater Video systems [95]. In addition, given the potentially longer time spend with the infield cables array, habituation to varying EMF levels is also a highly relevant research route. However, Hutchison et al 2020 hypothesize that elasmobranchs are less prone to habituation to SPC EMFs as these are highly variable in frequency distribution, exposure frequency and duration which makes them less predictable [47]. Thus in habituation research this variability should be carefully considered in the research design.
3. | **Interaction with migratory behaviour** Migratory species as *Mustelus asterias* and *Galeorhinus galeus* with alongshore migration routes during spring and autumn [61], [73], [74] have a high probability of exposure. Other more resident species, such as *Scyliorhinus canicula,* will have a lower probability of encountering DC SPC alongshore. The biggest risk to date lies in the uncertainty of the severity of impact from SPC on migratory behaviour and can consequently range from low (short duration) to high (strong deviation off course or delays). Long delays and deviation of course could result in failure to meet the migration goal in time or increased changes of predation due to a changed swimming depth. Research efforts could be focussed on reducing the uncertainties in the NOEL for the long-distance migratory species. This can be done for example done through tagging studies with a sensor that combines behavioural parameters (heart stroke volume, temperature, activity, tilt) and environmental parameters (salinity, depth) with a magnetometer. As long-term migration species are believed to use the earth magnetic field (DC), research on migration could be focused on DC cables which interact with the earth magnetic field.

**Figure 6.**
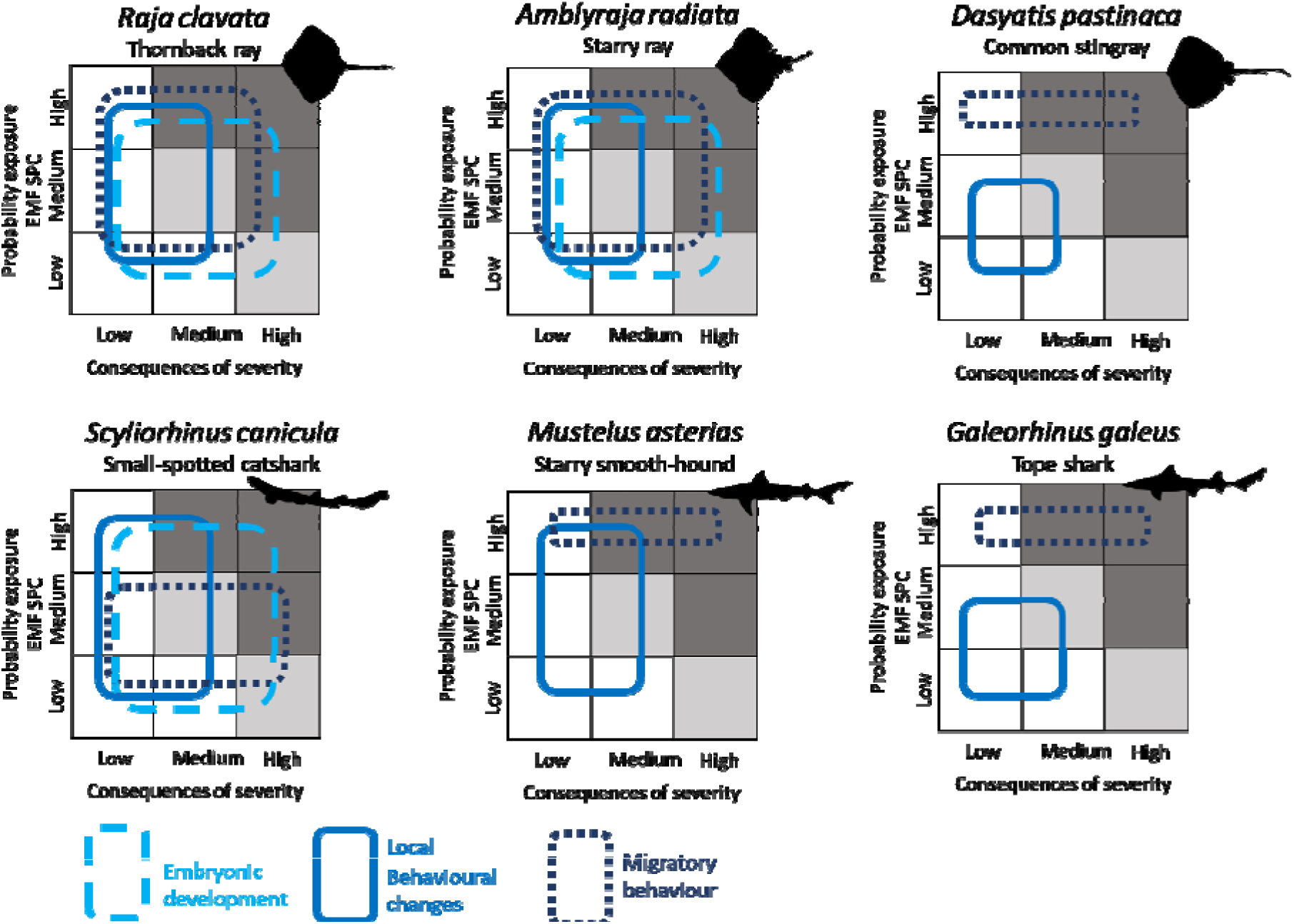
Risk assessment matrices for species with different ecology showing the range of risk for embryonic development, local behavioural changes and migratory behaviour in relation to electromagnetic fields exposure resulting from subsea power cables. The risk is categorised ranging from no to severe concern by assessing individual probability exposure to EMFs from SPC (y-axis) versus the severity of consequence (x-axis) (following Altenback, 1996). The resulting risk range differs depending on the biology and ecology of different species groups, e.g. rays, sharks or skates, resident or migratory, and oviparous or ovoviviparous/viviparous species. The white squares indicate no risk, the lightly shaded squares indicate minor risk and the dark shaded squares indicate high risk.

The risks discussed for the different behavioural categories describe single effects. A combination of effects under varying exposure levels, at different life stages and during a range of ecological activities could increase the probability of exposure and the severity of the effect. For example, when attraction of females near egg-laying sites is combined with impacts on embryogenesis once eggs are laid then the risks might be classified as more severe.

### 6.2. Validity of EMF exposure modelling approach for other countries

Whilst this study has focussed on the Dutch Continental Shelf as a case study of our approach, it is applicable in all seas with OWF and SPC expansion. In particular for Belgium, United Kingdom, German and Danish North Sea OWFs, as the species distribution range for the species discussed above overlap and migration for some species occur at transboundary scales. Moreover, our approach can also be applied conceptually for species with similar ecological traits. What is important to consider is that SPC will generate different EMFs levels in other countries if regulation results in different specifications, e.g. in cable design, burial depth and frequency on which AC cables operate, all of which will influence the magnetic field. Most notably, on the Dutch Continental Shelf bundled bipolar DC cables are the norm but in many countries the plus and minus cores are laid separately, with up to 250 m distance between them. This will result in elevated magnetic fields of up to 300 µT which is six times the earth magnetic field (∼50 µT on the Dutch Continental Shelf) and almost 2.5 times the maximum levels modelled for the Dutch situation in this study. Different cable specifications resulting in different EMFs can lead to a changed risk profile. In most countries there is no minimum burial depth or it is not possible to bury to a required depth due to rocky substrate or highly morpho-dynamic areas resulting in variable burial depths. Although these regional differences influence the level, characteristics and spatiotemporal fluctuations of EMFs exposure, the modelling approach presented here can be applied in other regions as well, provided the different cable specification and regional difference are taken into consideration.

The findings from the research priorities resulting from the ERA should provide valuable insights and support for the development of mitigation strategies or choices with respect to precautionary approaches pending upcoming studies and insights. In the case of observing an effect on embryogenesis, it is likely that rerouting the SPC would be the most suitable mitigation option, considering the localized nature of egg laying sites for benthic elasmobranchs. On the other hand, if the choice for egg-laying sites is related to habitat features associated with SPCs and OWFs this mitigation option should be reconsidered, again showing the need for dedicated research. If evidence for effects on migration is found, investigating SPC corridors (combining multiple cable routes) could be the most appropriate approach to minimize the risk of encounters [52].

### 6.3. Conclusion

The offshore energy transition is going full steam ahead across the marine environment, highlighting the importance to address the risks of EMFs from the huge expansion SPC on benthic elasmobranchs. Filling the knowledge gaps with the highest priority will either retire the risk or support policy decisions to prevent significant ecological impact through mitigation measures. As elasmobranch populations are already under the influence of many anthropogenic stressors, conservation or mitigation actions focussing on reducing impacts from EMFs should be properly considered in relation to other stressors as fishing, shipping and sand mining. The ERA approach set out here is a useful step forward in understanding research priorities for EMF.

## Acknowledgments

This work was funded by the Dutch Research Council NWO [grant number ENPPS.TA.019.005 (Elasmopower)]. We acknowledge the ElasmoPower steering group for their advice and review. We extend our gratitude to Jochem Boersma, Jelle Tams and Jobber Bekkers from consultancy firm Witteveen+Bos and Nick Schilder from TNO for their insights in the modelling of magnetic fields. We thank Katinka Bleeker and Jurgen Batsleer for providing the Frisbe and DATRAS database data and Brent Goossens for his assistance in visualising the data. Lastly we thank the Offshore Wind Parks owners Blauwwind, Eneco, Shell and Gemini and transmission system operator TenneT for providing data on the cables systems included in the modelling.

## Author Statement

AH and TM developed the risk assessment approach and designed the outline. AH wrote the first version of the manuscript and conducted the modelling assisted by those mentioned in the acknowledgements. All authors reviewed and commented on versions of the manuscript before submission.

## Supplementary materials

### Supplementary materials 1 - Description model

The calculations presented here are carried out and visualized using a model developed based on the Biot-Savart law. The Biot-Savart law is expressed as follows:

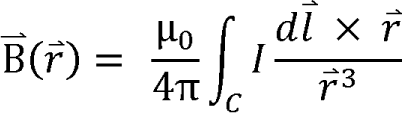

The magnetic field (B⃑) is dependent on the magnetic permeability (µ_0_), the amount of current (*I*) that flows through the cable, the geometry or direction of the cable (*dl⃑*) and the distance to the point at which the magnetic field is computed (*dr⃑*). When the distance between the cable and the point at which the magnetic field is to be computed is much smaller than the cable length, then the above equation can be simplified to the following:

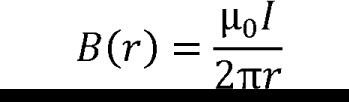

The above equation shows that the magnetic field is directly proportional to current flowing through the cable and inverse proportional to the distance r. The above equation computes the magnetic field for a single cable. In case of DC and AC cables there are multiple cables with different phases that need to be accounted for. Using the method shown by the National Grid of the United Kingdom on the Electric and magnetic fields and health (EMFS) webpage, the total magnetic field at a certain point can be computed as a result of the different phases of the different cables.

Source: How to calculate the magnetic field from a three-phase circuit’. EMFS, maintained by National Grid, https://www.emfs.info/wp-content/uploads/2014/07/Howtocalculatethemagneticfieldfromathree.pdf, accessed on 11-08-2023

**Supplementary Table 1.**
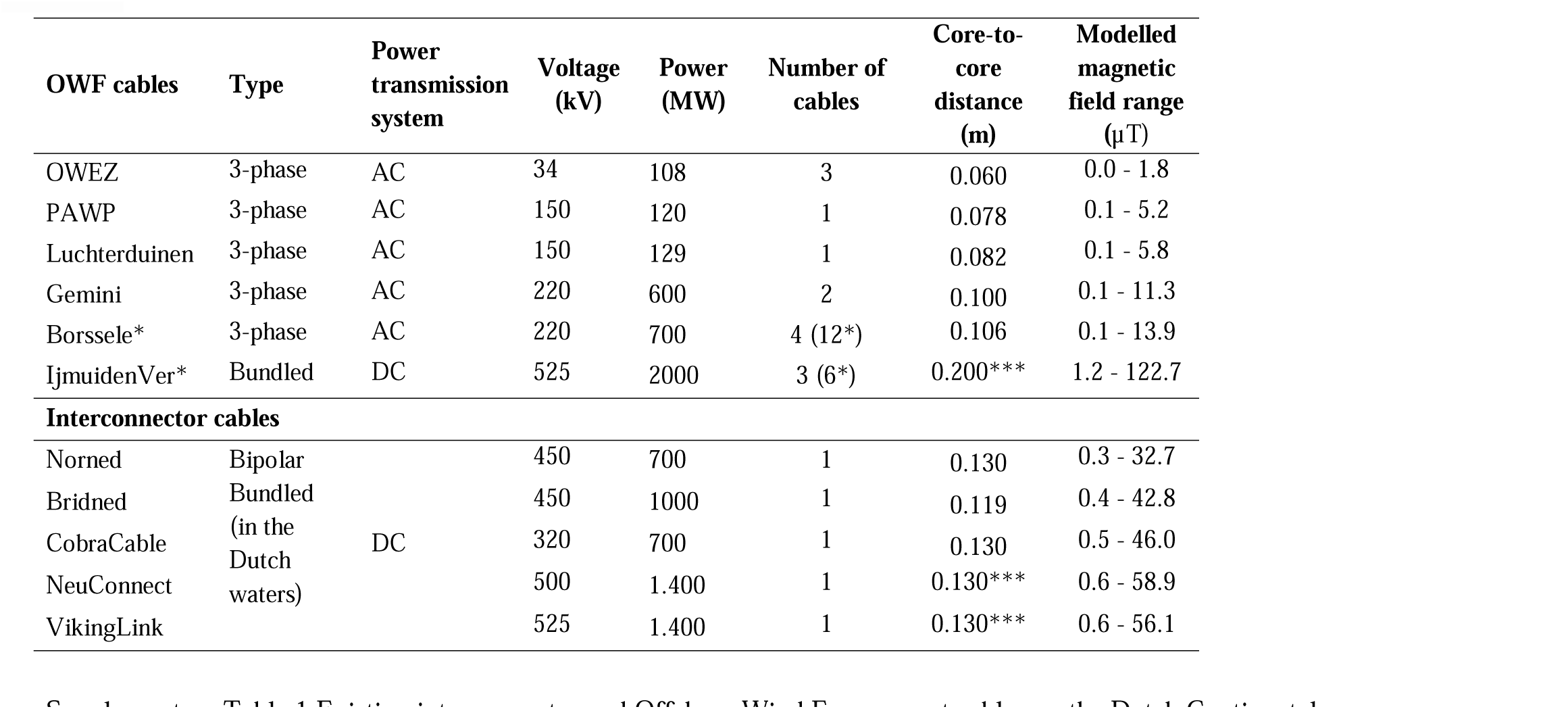
Existing interconnector and Offshore Wind Farm export cables on the Dutch Continental Shelf. * The design strengths for Borssele are the same for the planned offshore wind parks Hollandse Kust (Zuid, Noord, West) and Ten Noorden van de Wadden and Ijmuiden Ver are the same for Nederwiek. ** Cables for IjmuidenVer are planned to be bundled depending on the technical availably of bundled 2GW cables. *** Core-to-core distances are assumed when no public information or cable owner information was available.

**Supplementary Table 2.** Impact zone calculated based on the 5 nT zone and the length of the subsea power cable on the Dutch Continental Shelf.

| Cable | Length within Dutch Continental Shelf (km) | Amount of cables | 5 nT zone (m) from the centre of the cable | Affected area (km<sup>2</sup>) |
| --- | --- | --- | --- | --- |
| Norned | 57 | 1 | 89 | 10.2 |
| BRITNED | 101 | 1 | 103 | 20.8 |
| Cobra cable | 98 | 1 | 106 | 20.8 |
| Neuconnect | 349 | 1 | 121 | 84.5 |
| Viking Link | 203 | 1 | 118 | 47.8 |
| Ijmuiden Ver (alpha) | 163 | 1 | 180 | 58.7 |
| Ijmuiden Ver (beta) | 146 | 1 | 180 | 52.6 |
| Ijmuiden Ver (gamma) | 152 | 1 | 180 | 54.7 |
| Nederwiek (alpha) | 208 | 1 | 180 | 74.9 |
| Nederwiek (beta) | 200 | 1 | 180 | 72.0 |
| Nederwiek (gamma) | 278 | 1 | 180 | 100.0 |
| OWEZ | 15 | 3 | 20.7 | 5.7 |
| PAWP | 27 | 1 | 71 | 3.9 |
| Luchterduinen | 25 | 1 | 74 | 3.7 |
| Gemini | 91 | 2 | 52 | 38 |
| Borssele | 67 | 4 | 57.5 | 61.6 |
| HK(z) | 42 | 4 | 57.5 | 38.8 |
| HK(n) | 33 | 2 | 57.5 | 15.1 |
| HK(w) | 64 | 4 | 57.5 | 59.4 |
| TNW | 82 | 2 | 57.5 | 37.9 |
| Total | 2401 | 34 | - | 861.1 |

